# B.1.617.3 SARS CoV-2 spike E156G/Δ157-158 mutations contribute to reduced neutralization sensitivity and increased infectivity

**DOI:** 10.1101/2021.10.04.463028

**Authors:** Tarun Mishra, Garima Joshi, Atul Kumar, Rishikesh Dalavi, Pankaj Pandey, Sanjeev Shukla, Ram Kumar Mishra, Ajit Chande

## Abstract

SARS CoV-2 variants raise significant concerns due to their ability to cause vaccine breakthrough infections. Here, we sequence-characterized the spike gene, isolated from a breakthrough infection, that corresponded to B.1.617.3 lineage. Delineating the functional impact of spike mutations using reporter pseudoviruses (PV) revealed that N-terminal domain (NTD)-specific E156G/Δ157-158 contributed to increased infectivity and reduced sensitivity to ChAdOx1 nCoV-19 vaccine (Covishield™)-elicited neutralizing antibodies. A six-nucleotide deletion (467-472) in the spike coding region introduced this change in the NTD. We confirmed the presence of E156G/Δ157-158 in the RT-PCR-positive cases concurrently screened, in addition to other circulating spike (S1) mutations like T19R, T95I, L452R, E484Q, and D614G. Notably, E156G/Δ157-158 was present in more than 85% of the sequences reported from the USA, UK, and India in August 2021. The spike PV bearing combination of E156G/Δ157-158 and L452R further promoted infectivity and conferred immune evasion. Additionally, increased cell-to-cell fusion was observed when spike harbored E156G/Δ157-158, L452R, and E484Q, suggesting a combinatorial effect of these mutations. Notwithstanding, the plasma from a recovered individual robustly inhibited mutant spike PV, indicating the increased breadth of neutralization post-recovery. Our data highlights the importance of spike NTD-specific changes in determining infectivity and immune escape of variants.

## Introduction

There was a substantial rise in the number of COVID-19 reported cases in India after March 2021, reaching more than 400000 cases per day ^1^. The extent of spread was attributed to fitness conferring mutations in the parental lineage B.1.617, leading to the emergence of sublineages like B.1.617.1, B.1.617.2, and B.1.617.3 of SARS CoV-2 ^2^. The delta variant belonging to B.1.617.2, detected in December 2020, dominated during the second wave in the country^3,4^. This emergence of variants coincided with the vaccination drive, prioritized for the frontline workers, older population with subsequent rollouts in high-risk groups and young adults. While the frontline workers mostly received both doses of ChAdOx1 nCoV-19 (Covishield™ in India) by March 2021, highly transmissible delta and delta plus variants displayed the ability to cause breakthrough infections, challenging the vaccine efficacy ^5^. Our community surveillance analysis also identified a few cases that were classified as vaccine breakthrough infections. Rigorous testing and isolation were carried out to prevent further spread, and it became pertinent to understand the potential of the pathogen targeting vaccinated individuals.

SARS CoV-2 entry is mediated by the interaction of its spike (S) protein on the virions with the human ACE2 receptor ^6^. Spike protein variants harboring L452R and E484Q mutations were reported to have contributed to the pathogenicity of the virus (**Ferreira et al., 2021; Michael Rajah et al., 2021**). These mutations are found in the critical receptor-binding domain that interacts with the ACE2 and is a target for neutralizing antibodies. Indeed, recent reports demonstrated diminished sensitivity of spike PV bearing L452R and E484Q to BNT162b2 mRNA vaccine-elicited antibodies but a lack of synergy between these two mutations in conferring the resistance ^7,9^.

While spike-focused first-generation vaccines designed based on the seeding variants have been shown to prevent symptomatic disease effectively ^10,11^, the ability of the delta variant, like other reported variants, was envisaged to be an attribute of spike protein that enabled the virus to evade vaccine-stimulated host defense ^12,13^. Despite the restricted tropism to ACE2 expressing cells, the spike appears tolerant to mutations that confer the ability to escape humoral immunity and, by extension, the resistance to antibody treatments ^14^. Therefore, to understand the molecular determinants in the spike protein responsible for reduced vaccine effectiveness, we cloned and sequence-characterized the spike gene from a vaccine breakthrough infection case. The spike nucleotide sequence analysis revealed a series of mutations that we functionally characterized using reporter PVs (Mishra et al. 2021). We observed that amino acid changes in the B.1.617.3 SARS CoV-2 spike NTD contribute to increased infectivity, resistance to vaccine-elicited polyclonal antibodies, and cell-to-cell fusion.

## Results

### Spike gene from breakthrough infection harbors a six-nucleotide deletion

During our surveillance study, we identified a previously uninfected, ChAdOx1 nCoV-19 (Covishield™) fully-vaccinated case that was infected 50 days after the second dose. In an effort to map the spike mutations that plausibly enabled the virus to dampen the host defense, we PCR amplified a full-length gene from the reverse transcribed RNA. Primer walking utilizing Sanger sequencing followed by analysis resulted in the contigs that were assembled, and mutations were scored to comprehend the origin of the spike protein variant (termed hereafter ICS-05). We first compared the spike sequence from the initial Wuhan isolate (Figure-1a; Supplementary figure-1). We found a total of eight changes; three were in NTD (N- terminal domain), four in RBD (receptor binding domain), and one on the S2 portion of ICS-05 spike (Figure-1a). Interestingly, we observed a six-nucleotide deletion that resulted in the loss of two amino acids at 157 and 158 positions and a change of Glutamic acid at 156 positions to glycine (E156G/Δ157-158) (Supplementary figure-1). This deletion we found in five of the total seven spike sequences isolated from the RT-PCR positive cases (Supplementary figure-1). Out of these five cases, two were fully vaccinated. When the ICS-05 spike sequence was aligned with the available spike sequences on GISAID^15^, it corresponded to the B.1.617 lineage, specifically with B.1.617.3 of the delta variant of concern (Figure-1b). Surfaced in the month of March 2021, the delta variant dominated the second wave in the country (Figure-1c) and was reported to have caused 25.3% of breakthrough infections ^5^. The E156G/∆157-158 mutation, first detected on 7^th^ August 2020, subsequently became 35% prevalent worldwide (Figure-1d), and by August 2021, it was found in more than 85% of reported sequences from the USA, UK, and India (Figure-1f). The E156G/∆157-158 mutation has been detected with high frequency (Figure-1e) in at least 157 countries and is found in multiple PANGO lineages (**Abdel Latif 2021; Elbe and Buckland-Merrett, 2017**). Given the higher prevalence worldwide and in the most affected countries (Figure-1d, 1f), we hypothesized a functional relevance of E156G/Δ157-158 mutations.

**Figure-1.**
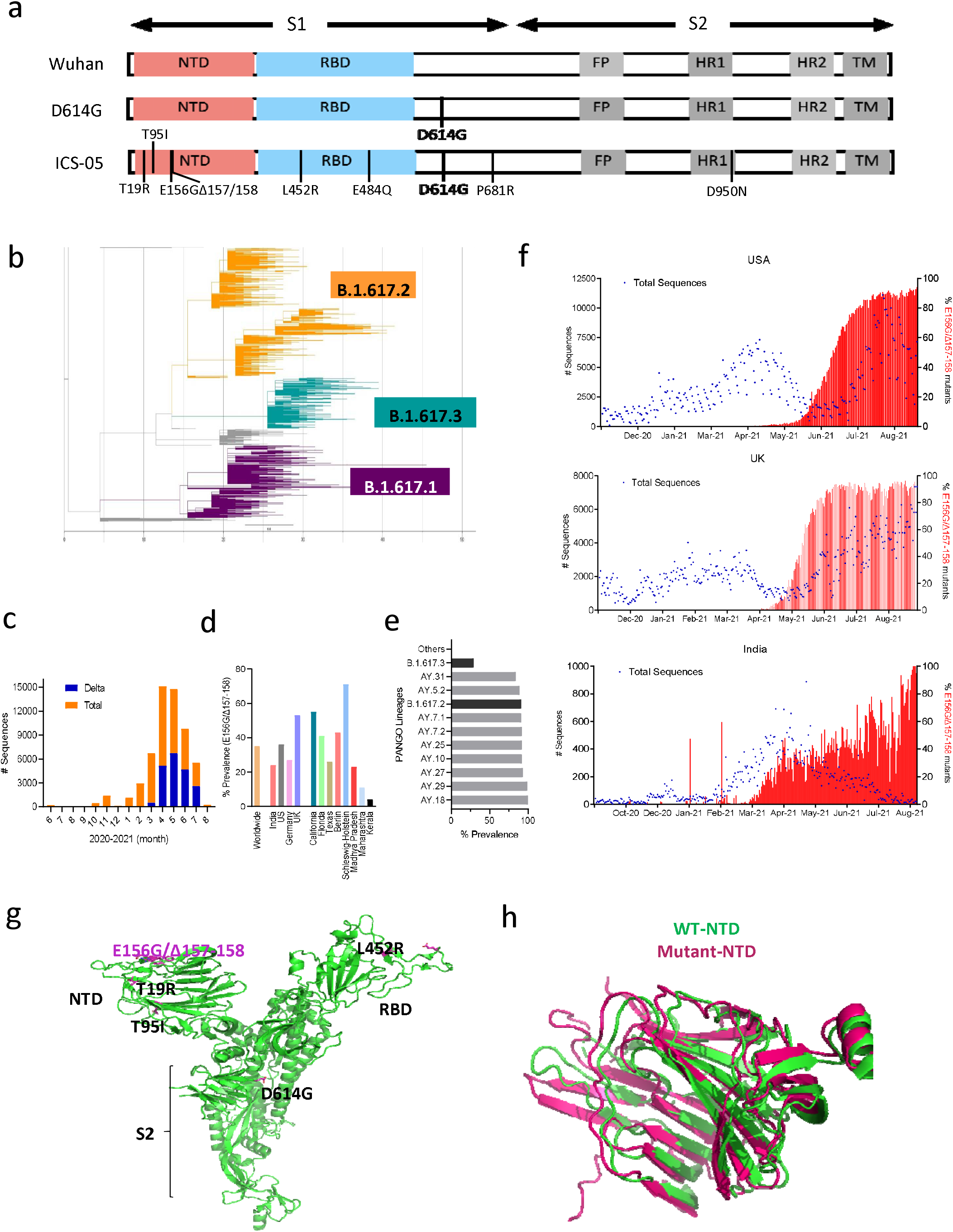
Sequence characterization, geographical prevalence, and structural significance of spike mutations. a. Schematics of spike protein representing the mutations in the S1 and S2 domains. Different regions of spike proteins are indicated: NTD (N-terminal domain), RBD (receptor binding domain), FP (fusion peptide) HR1 & 2 (heat repeat 1 and 2), TM (transmembrane region). b. SARS-CoV-2 complete sequences (2788) from GISAID were checked for quality on Nextstrain. The maximum likelihood tree was interpreted using Fig-tree. The multiple alignments were performed using nextstrain pipeline on these 2788 complete genomic sequences of SARS CoV-2. Branch lengths denoted the number of nucleotide substitutions from the root of the tree. The key lineages were colored eggplant B.1.617.1, yellow B.1.617.2, turquoise B.1.617.3. c. Bar-graph indicates the frequency of Delta variants during the second wave in India. The yellow color bar shows the total number of SARS COV-2 sequences submitted each month (denoted on the x-axis), and the blue bar represents the number of sequences among total sequences submitted between the period of June 2020- August 2021. The data was obtained from the GISAID SARS CoV-2 database (outbreak.info). d. Bar-graph represents the prevalence of E156G/∆157-158 mutation in the indicated countries/state/worldwide. e. Occurrence of E156G/∆157-158 in the PANGO lineages (https://cov-lineages.org/index.html). f. The numbers of sequences carrying E156G/∆157-158 were reported by indicated countries. The left y-axis represents the total number (#) of SARS CoV-2 genome sequences (blue dots), while the right y-axis denotes the percentage of E156G/∆157-158 occurrence (red bars). The X-axis represents the months of sequence submission. g. mutations found in the spike ICS-05 are shown in the sticks (magenta) in Spike protomer (green, pdbid:7DF3). h. Superimposition of WT NTD (green, pdbid:7DF3) and E156G/∆157-158 NTD (magenta) of the spike protein.

We first examined the spike protein in the structural context of E156G/Δ157-158 for clues regarding the alteration of epitopes. Interestingly, mapping of E156G/Δ157-158 on the structure of wild-type spike protein shows that the mutated region is surface exposed, which might be a good target for antibodies (Figure-1g). Further to understand the effect of these mutations on spike protein structure, we predicted the structure of NTD bearing E156G/Δ157-158 using the AlphaFold^18^. The resultant model of a mutant-spike protein does not show any significant changes in the NTD of spike protein (Figure-1h), suggesting resistance to neutralizing antibodies may not be attributed to structural changes.

### E156G/Δ157-158 contributed to attenuated susceptibility to neutralization and promoted infectivity

To appraise the exact potential of the NTD bearing E156G/Δ157-158 and changes in the region important for the receptor binding, we introduced the indicated mutations on the reference D614G spike by site-directed mutagenesis (Figure-2a). Next, we assessed the effects of these spike mutations on the infectivity of PV following our earlier report ^19^. The lentiviral spike PVs carried a luciferase gene, and the values were represented after normalizing to the milli units of reverse transcriptase (RT mU). In agreement with previous findings (**Ferreira et al., 2021**), while the RBD-specific mutation E484Q did not significantly confer infectivity advantage to the spike particles, the L452R mutation increased the infectivity more than two-fold in these conditions (Figure-2b).

**Figure-2.**
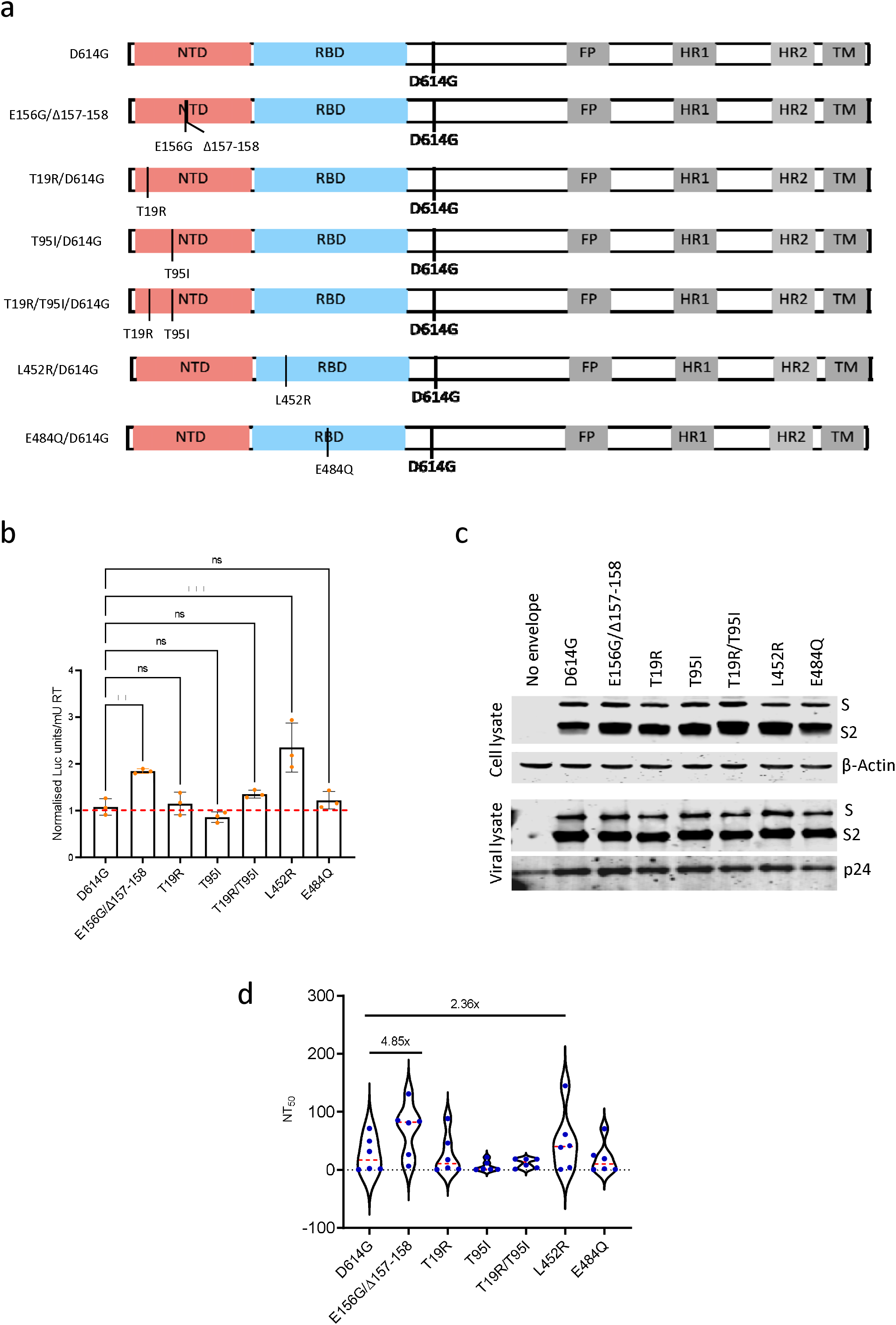
Infectivity and neutralization of spike pseudoparticles. a. Schematics of the spike mutants generated to study the effect of ICS-05-specific mutations. Amino acid positions are represented with respect to the Wuhan HU-1 sequence (NC_045512). b. Infectivity profiles of the indicated spike mutant-pseudotyped lentiviruses. The infectivity was normalized to the D614G pseudotyped lentiviral particles. The data represent the mean of three replicates, and the significance was measured by one-way ANOVA multiple comparison test to analyze the difference between the groups, n=3. *p<0.05, **p<0.01, ***p<0.001, ns=non-significant. c. Western blots indicate the relative expression of the indicated spike proteins bearing mutations from- the producer cell lysates and -the viral lysates. Beta-actin and p24 served as loading controls for cell lysates and viral lysates, respectively. d. The susceptibility of each spike mutant PV to neutralization by antibodies in the plasma obtained from vaccinated, test-negative individuals. The dotted red line represents the median response of each spike PV. The fold difference in response to neutralizing plasma was measured compared to the reference D614G mutant spike PV (n=6). The statistical significance was calculated by the Wilcoxon Signed Rank test, two-tailed, non-parametric.

Interestingly, the spike bearing E156G/Δ157-158 mutation was almost equally infectious compared to the L452R mutant, indicating a potential change in the NTD that contributed to increased PV infectivity. The remaining mutations examined (T19R, T95I, T19R/T95I) did not significantly confer infectivity advantage (Figure-2b). Western blotting from the producer cell lysates and purified virions indicated that all these spike protein mutants were expressed, and there was no noticeable effect on the spike processing or virion incorporation (Figure-2c).

Next, we examined the susceptibility to neutralization of indicated spike PV to vaccine-elicited plasma polyclonal antibodies from test-negative individuals. With the D614G as a reference, the NT_50_ values obtained showed that spike PV carrying E156G/Δ157-158 mutation was 4.85-fold less susceptible to vaccine-elicited polyclonal antibodies (Figure-2d), indicating the role of this mutation in escaping the vaccine-elicited antiviral immunity in addition to promoting virion infectivity. The other indicated mutations did not confer noticeable resistance in these conditions except the L452R mutant bearing PV that required a 2.36-fold higher plasma for neutralization, consistent with the previous findings ^7^ (Figure-2b and 2c). Altogether, these results suggest the contribution of the NTD-specific mutations in conferring resistance to neutralization and promotion of infectivity.

### E156G/Δ157-158 and L452R additively effects immune escape and increased infectivity

Plasma polyclonal antibodies post-recovery were capable of neutralizing the ICS-05 spike PV despite the presence of mutations (Supplementary Figure-2), suggestive of the increased breadth of neutralization. For delineating key events shaping the escape from polyclonal neutralizing responses, we compared the effect of combinations of these select mutants (Figure 3a) on PV neutralization by vaccine-elicited antibodies. Specifically, we asked if different mutations acted in synergy in the antibody evasion process. The ability of E484Q and L452R to evade BNT162b2 Pfizer mRNA vaccine-elicited antibodies has been established recently ^14,20,21^. We combined the NTD-specific change E156G/Δ157-158 with E484Q and L452R (Figure-3a) and performed the infectivity and neutralization assays. Irrespective of the background (E484Q or the L452R), NTD-specific mutation E156G/Δ157-158 increased the infectivity ∼4-fold for the spike pseudotyped lentiviral particles (Figure-3b). Western blotting experiments revealed that these mutant spikes were enriched almost equally in the virions; however, the expression of E156G/Δ157-158/ L452R/E484Q and ICS-05 spike in the cell lysates was not comparable with other counterparts (Figure-3c). The expression discrepancy might be attributed to a specific condition that is needed to solubilize these mutants. Nevertheless, we see that the virion incorporated fraction was similar for all mutants, including ICS-05, indicating that the associated phenotypes are unlikely affected. Notably, the reduced susceptibility to neutralization observed for the ICS-05 spike (11-fold) was mostly conferred by a combination spike-mutant that harbored E156G/Δ157-158 and L452R (7-fold less susceptible to neutralization) (Figure-3d).

**Figure-3.**
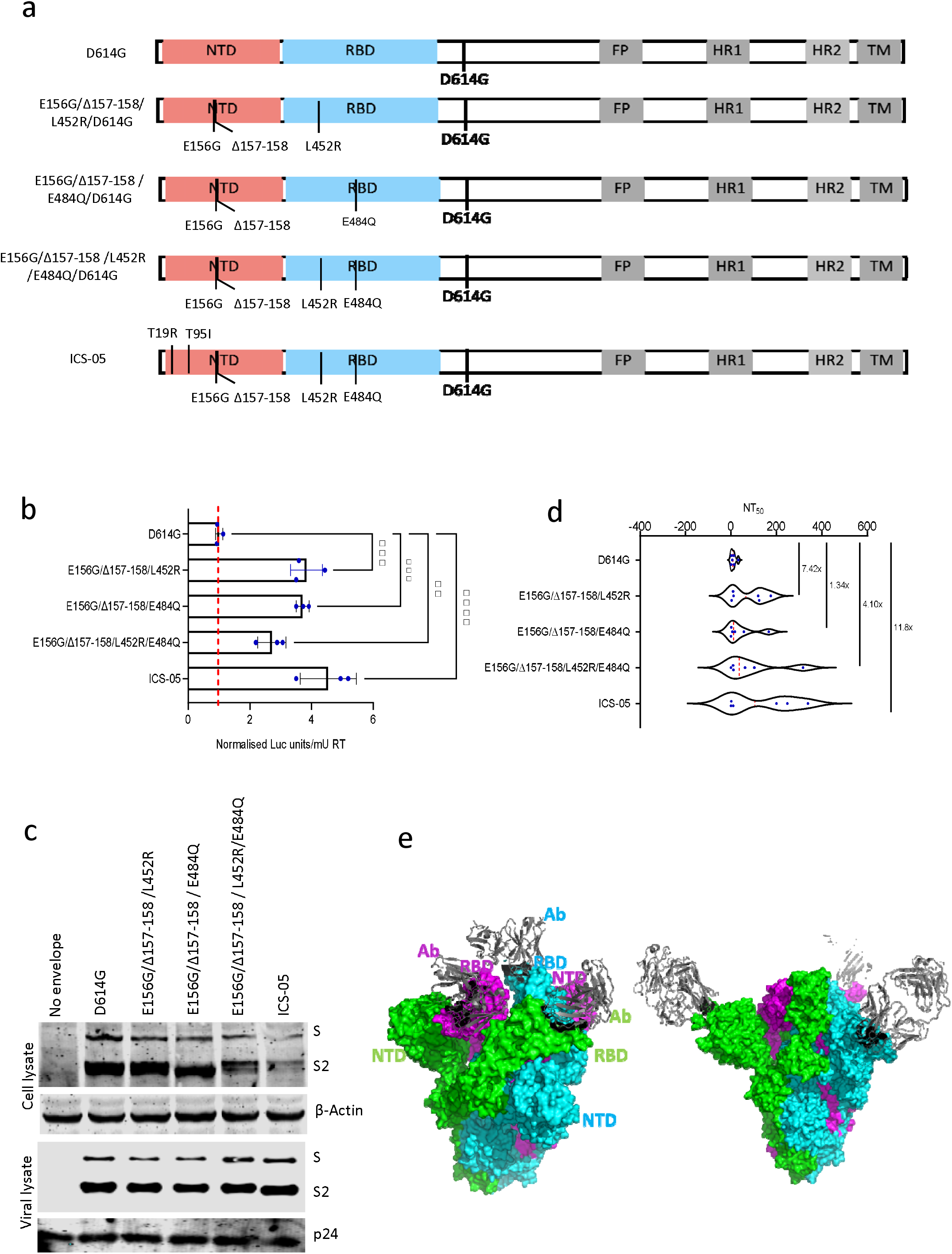
Infectivity and neutralization of spike pseudoparticles and structural analysis. a. Schematics of the spike mutants generated to check the combined effects of mutations. Amino acid positions are represented with respect to the Wuhan HU-1 sequence (NC_045512). b. Infectivity profiles of the indicated spike mutant-pseudotyped lentiviruses. The infectivity was normalized to the D614G pseudotyped lentiviral particles. The data represent the mean of three replicates, and the significance was measured by one way ANOVA multiple comparison test to analyze the difference between the groups, n=3. *p<0.05, **p<0.01, ***p<0.001, ****p<0.0001. c. Western blots indicate the relative expression of the indicated spike proteins bearing mutations from- the producer cell lysates and -the viral lysates. Beta-actin and p24 served as loading controls for cell lysates and viral lysates, respectively. d. The susceptibility of each spike mutant PV to neutralization by the antibodies in the plasma obtained from the vaccinated, test-negative individuals. The dotted red line represents the median response of each spike PV. The fold difference in response to neutralizing plasma was measured compared to the reference D614G mutant spike PV (n=6). The statistical significance was calculated by Wilcoxon Signed rank test, two-tailed, non-parametric. e. The complex structure of RBD-specific antibody bound with a trimer of spike proteins (pdbid:6XEY). The surface of spike protein protomers are shown in green, cyan, and magenta, antibody (grey) bound to RBD of spike protein are shown as a cartoon (left panel). The complex structure of NTD specific antibody bound with a trimer of spike proteins (pdbid:7C2L). The surface of spike protein protomers is shown in green, cyan, and magenta, antibody (grey) bound to NTD of spike protein is shown as a cartoon (right panel).

To further understand the reduced susceptibility to neutralization upon alterations in the NTD of the spike, we explored known complex structures of spike with antibodies. There are two major classes of antibodies; one binds with the NTD region, whereas another binds with the RBD domain of spike protein. The structure of antibodies bound to RBD and NTD domain of spike reveals that the neutralization escaping mutations described in this study are present at the interface of antibody and NTD and/or RBD domain of spike protein (Figure-3e). This observation is consistent with our neutralization assay results which shows that mutations in these regions affect PV sensitivity to neutralization. Altogether, we observed that these mutations in NTD, particularly E156G/Δ157-158, cooperated with the seeding changes in the RBD, like L452R, for neutralizing antibody escape.

### Cell-to-cell fusion is enhanced by the NTD and RBD specific amino acid changes

The pathogenicity of SARS CoV-2 is also attributed to spike-dictated formation of syncytia, a phenotype characterized by cells with abnormal morphology and frequent multinucleation. This feature has been more pronounced with delta variants ^22,23^. Cell-to-cell fusion also offers an avenue for evasion of humoral responses and rapid viral dissemination ^22,24^. Our analysis revealed that E156G/Δ157-158 together with L452R conferred most of the resistance to antiviral immunity elicited by vaccination. We next asked if these amino acid changes also promoted cell-to-cell fusion leading to syncytium formation. We designed assays that revealed the ability of the cells expressing indicated spike mutants (Red) to fuse with bystander ACE2 expressing cells (Green). While E156G/Δ157-158 alone had an indiscernible effect on syncytium formation, the area after cell-to-cell fusion was markedly increased when the cells expressing D614G spike bearing E156G/Δ157-158, L452R, and E484Q mutations were co-cultured with ACE2-positive cells (Figure-4a and 4b). These results indicated the increased ability of the ICS-05 spike is conferred by NTD- and RBD-specific changes that acted in concert to promote syncytium formation.

**Figure-4.**
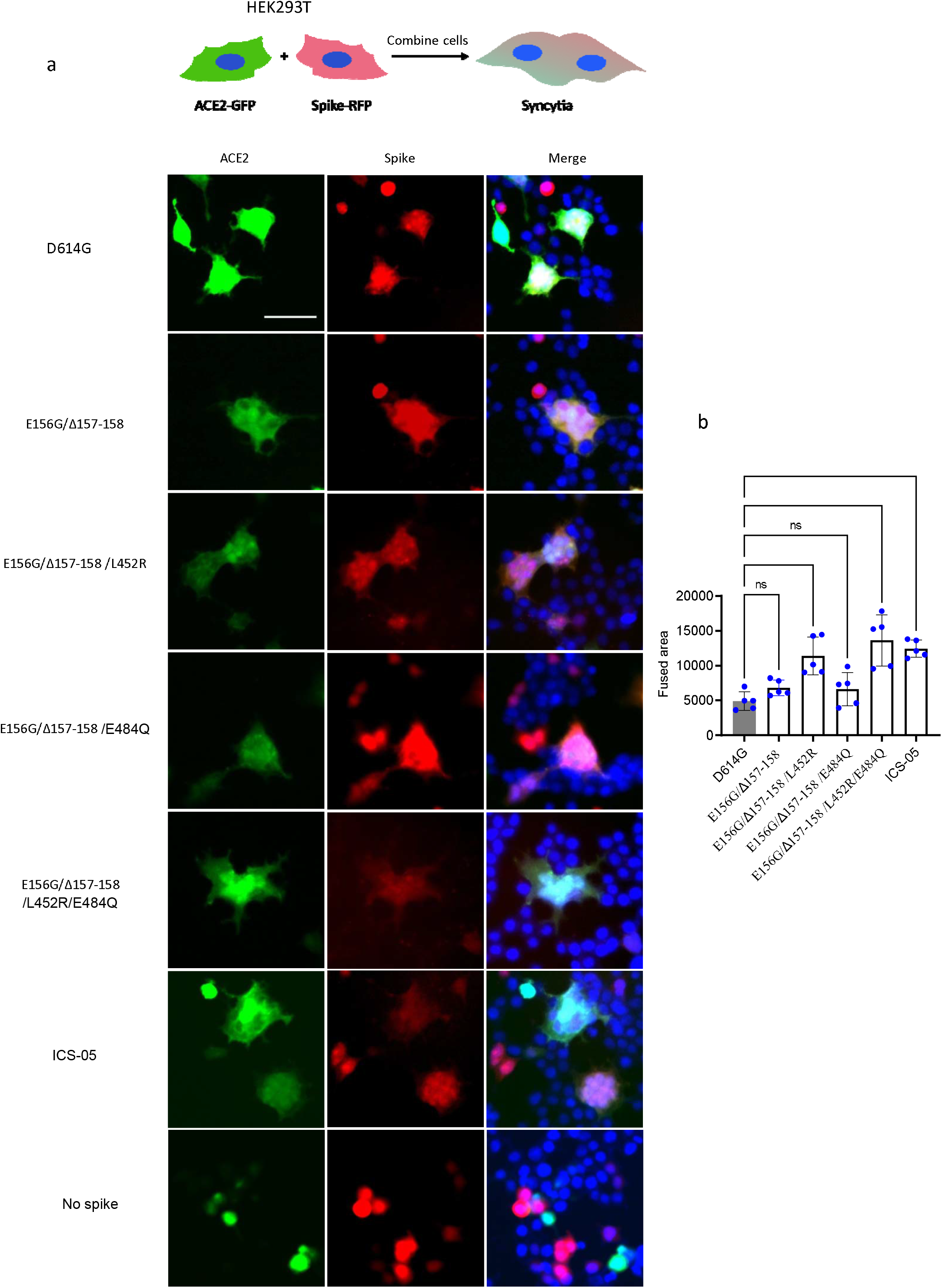
SARS CoV-2 spike mutants form syncytia in ACE2 expressing cells. a. The HEK293T ACE2 transfected cells (green) were mixed with indicated spike mutants transfected cells (red) displaying syncytia formation. The cells with ACE2 expression mixed with the control cells with no spike expression served as control. Scale 50μm. b. Quantification of the cell-cell fusion by measuring the fused area for five different fields from three replicates. The bar graph represents mean±SD. Statistical significance was calculated by one-way ANOVA.

## Discussion

Little is known about the functional consequences of NTD-specific changes in the spike of emerging SARS CoV-2 variants. In contrast, RBD-specific changes in the spike have been considered a defining feature that confers fitness to the virus. Here, we demonstrated that molecular features of the spike (like L452R and E484Q), already known to confer fitness ^7,25,26^, cooperates with alterations in the NTD to enhance spike function further. The now widespread NTD-specific mutation E156G/∆157-158 likely conferred an evolutionary advantage and might underly vaccine breakthroughs found common during the delta outbreaks ^5^. One of Delta’s siblings, a variant called Lambda that, like Delta, also carries changes in the spike NTD, in addition to L452Q, and these alterations have been linked to the virus’ higher infectivity and immune evasion ability ^27,28^. In agreement with these observations, we found that the ICS-05 spike carried mutations in the NTD coding sequence and that these changes indeed acted in concert in evading antiviral immunity elicited by the vaccine and contributed to increased infectivity. Furthermore, consistent with the previous reports, we also observed ACE2 expressing cells forming larger syncytia when mixed with spike-expressing cells, particularly ICS-05 spike ^8,23,29^. It has been found that the extent of syncytia formation in SARS CoV-2 infected patients’ lungs positively correlates with disease severity and higher mortality ^22,24^. Our observations that E156G/Δ157-158, L452R/E484Q mutations bearing spike induced large syncytia formation, almost equivalent to that of ICS-05 spike, exemplifies NTD and RBD-specific changes that together can further promote cell-cell fusion.

Our analysis identified a six-nucleotide deletion in the ICS-05 spike isolate from a breakthrough infection case, concurrently found in more cases regardless of the vaccination status. The prevalence of this deletion in the sequences reported worldwide and in the most affected countries suggests its impact on virus transmission. Out of the seven cases that were screened, two were fully vaccinated and carried spike E156G/Δ157-158 mutations. We used plasma collected from frontline workers who were vaccinated on the same day/place and received two doses at identical intervals. The variability observed in the neutralization profiles using PVs, however, was expected as it basically reflects differential antibody responses between various individuals.

One of the limitations was that we did not have access to the plasma from before infection to appreciate the vaccine efficacy for the subject case. Furthermore, undoubtedly, the sample size would have further strengthened the study. However, we could not have had more samples that were matched for geographical location, the time intervals between the two doses (later on changed) given the limited numbers of vaccinated individuals then. Within these limitations, our results have important implications suggesting the possibility that NTD specific changes, like the ones reported here, might be driving the spread of variants.

We have confirmed that the L452R mutation in the RBD, prevalent in the Delta, Lambda, and epsilon variants, contributes to enhancing infectivity ^4,21,30^. The Delta and Lambda variants under VOI and VOC, respectively, do not alone rely on the higher infectivity for spread but also showed immune evasion from the vaccine-induced antibodies ^7,14,20^. In contrast, the Epsilon variant is now removed from the VOC/VOI list because of its low spreading capacity, although being more infectious than the spike bearing D614G. This suggests that higher infectivity alone is not sufficient for the spread. As shown here, the combined effect of E156G/Δ157-158 and L452R mutations that maintain higher infectivity and reduced neutralization susceptibility may underlie a particular variant’s dominance.

The growing evidence that NTD-specific mutations are modulating antigenicity requires immediate attention. Some NTD targeting mAbs, isolated from recovered patients, have been structurally characterized ^31,32^. Recent studies functionally evaluated the mutations in NTD of spike protein that help the virus transmit better and escape from neutralizing antibodies ^30,32,33^. As observed from the sequencing data on GISAID, several sequences belonging to emerging variants are found with a deletion/mutation in the NTD region of the spike. Multiple beta variants contain three amino acid deletion LAL242-244 NTD ^25,30^, which does not change the neutralization effect of antibodies, but specific antibodies like 4A8 targeting the NTD are shown not to neutralize the virus bearing these changes in the spike ^25,34^. Most recently, the lambda variant possessing the RSYLTPGD246-253N mutation in NTD have been shown to confer resistance to vaccine-elicited antibodies, particularly antibodies targeting the NTD “supersite” ^28,32,33,35,36^. In agreement with these reports, we show that, under immune pressure, the SARS CoV-2 spike can use mutations in NTD, along with the RBD, to escape polyclonal neutralizing responses.

Vaccine rollout programs coincided with the emergence of genetically distinct SARS CoV-2 variants. Right now, the extent to which spike mutations will enable these variants to evade vaccine-elicited immunity remains to be determined. Thus, more such studies aimed at understanding phenotypic impacts of globally circulating spike amino acid changes can inform the development of next-generation vaccines and mAbs.

## Methods

### Ethics statement

The Institute Ethics committee approved this study. All the samples were collected with due consent from the donors.

### Plasmids

The Plasmid expressing SARS CoV-2 Spike protein (Wuhan isolate) was procured from Addgene with 19 nineteen amino acids deletion at C- terminal that enables efficient lentiviral packaging. The pcDNA 3.1 bs (-) spike D614G mutant was generated by site directed mutagenesis first^19^. All other plasmids expressing spike protein mutants (pcDNA 3.1 bs(-) T19R, pcDNA 3.1 bs(-) T95I, pcDNA 3.1 bs(-) E156G/Δ157-158, pcDNA 3.1 bs(-) L452R, pcDNA 3.1 bs(-) E484Q, pcDNA 3.1 bs(-) E156G/Δ157-158/L452R, pcDNA 3.1 bs(-) E156G/Δ157-158/E484Q, pcDNA 3.1 bs(-) E156G/Δ157-158/L452R/E484Q, pcDNA 3.1 bs(-) ICS-05 were generated using site directed mutagenesis by PCR using pcDNA 3.1 bs(-) spike D614G plasmid as template. The following primers S: D614G forward; 5′gtgctgtaccagggcgtgaattgcacc3′ reverse; 5′ggtgcaattcacgccctggtacagcac3′ ; S: T19R forward; 5′tctggtctcgtctcagtgcgtgaacctgagaactagaacccagctgcctc3′ reverse; 5′ctagcagcagctgccgcagga3′; S: T95I forward; 5′gcgtgtacttcgcctccattgagaagagcaacatcatc3′ reverse; 5′gatgatgttgctcttctcaatggaggcgaagtacacgc3′ ; S: E156G/Δ157-158 forward; 5′aaggtgcaattgttggcggagctgtacacgccgctctccatccaggact3′ reverse; 5′agtcctggatggagagcggcgtgtacagctccgccaacaattgcacctt 3′ ; S: L452R forward; 5′gcaactacaattaccggtaccgcctgttccg3′ reverse 5′cggaacaggcggtaccggtaattgtagttgc3′; S: E484Q forward; 5′ccatgcaatggagtgcagggcttcaactgct reverse; 5′agcagttgaagccctgcactccattgcatgg3′ were used to generate the above mention plasmids. We use the term mutation to indicate an amino acid change with respect to Wuhan-Hu-1 reference sequence (NC_045512). All the constructs were sequence verified for the reported mutations. A list of plasmids and the relevant information is tabulated in Supplementary table-I.

### Cell Culture and reagents

HEK293T cells (ECACC) were grown in DMEM medium supplemented with 10% FBS, 2mM glutamine, and 1% penicillin-streptomycin. ACE2+ cells were generated by Lentiviral transduction of HEK293T cells and were selected on the hygromycin. Reagents details are provided in the supplementary table-II.

### Pseudovirus production

HEK293T cells (3×10^6^ cells) were seeded in a 10cm^2^ plate 24h before transfection for the spike-pseudotyped lentiviruses production. The cells were co-transfected, with pScalps Zsgreen Luciferase (8μg), psPAX2 (6μg), pcDNA 3.1 bs(-) N protein-encoding plasmid (2ug) and either 2μg of the parental spike (D614G) or its derivatives plasmids, by calcium phosphate transfection method ^19^. The cells were replenished with a fresh medium after 16h of transfection. The supernatant containing the viral particles was collected 48h post-transfection. The supernatant was centrifuged at 300g for five minutes and passed through a 0.22μm filter to remove cell debris. The viruses were quantified using an SGPERT assay ^37,38^. To check the virion incorporation of spike protein, the supernatant containing viral particles was overlayed on top of sucrose (20%) cushion and centrifuged at 100,000×g for 2h at 4°C. After centrifugation, supernatant was removed completely, and the virus pellet was resuspended in Laemmli buffer containing 10mM TCEP as a reducing agent.

### Transduction and infectivity measurement

All the infectivity experiments were performed in 96-well plate formats with 40-60% target cell confluency. For infectivity experiments, HEK293T ACE2+ cells were seeded 24h before transduction. The transduction was done in quadruplicate with dilutions (Undiluted, 1:5, 1:25, 1:125) of pseudotyped virus preparation as described earlier ^19,39,40^. The cells incubated with a growth medium with heat-inactivated FBS alone were considered as the control. The level of transduction was quantified using Luciferase assay.

### Luciferase assay

The luciferase assay was performed to quantify the level of transduction by spike variants lentiviruses. For this, the growth medium was removed from each well, and cells were washed with 1XPBS. Further, 100μl of lysis buffer was added to each well to lyse transduced cells at room temperature for 20 minutes. 50μl of lysate was transferred to a white 96-well plate and finally mixed with the 50μl of the substrate solution, and enzyme activity was measured using spectramaxi3X (Molecular Devices, USA).

### Collection of plasma and pseudovirus neutralization

For obtaining plasma, 5ml of venous blood sample was collected in EDTA containing vials and mixed gently by inverting multiple times to avoid coagulation. Further, the tubes were centrifuged at 200g for 15 minutes at 4°C in a swing-out bucket. The plasma layer above the PBMCs was collected as a source of antibodies. All the plasma samples used in the study were heat-inactivated (56°C, 30 minutes) to disrupt complement components. The plasma of the individual, who got two doses of Covishield and became COVID-19 positive, was collected after the RT-PCR negative report post-recovery. The plasma from test-negative vaccinated individuals who got both doses of the Covishield vaccine was collected after 30 days of vaccination.

To check the neutralization potential of antibodies, present in the plasma of vaccinated (Covishield) and a vaccinated individual post-COVID-19 recovery, the various amount (μl) of plasma (10, 1, 0.1, 0.01, 0.001, 0.0001, 0.00001, 0.000001) were incubated with equivalent Spike PV for 20 minutes at room temperature before challenging the HEK293T ACE2+ cells. Transduction efficiency was measured after 48h by luciferase assay as described.

### Cell-to-cell fusion assay

For studying fusion of ACE2 expressing cells with spike expressing cells, we seeded HEK293T cells in 24 well plates at 70-80% confluency and co-transfected separately each well with Tag-RFP 657 (50ng) along with spike protein expressing vector (500ng) harboring indicated mutations. Separately, HEK293T cells were co-transfected with pEGFPN1 (50ng) and ACE2 expressing vector (500ng). After 10h of transfection, cells were trypsinized, mixed at 1:1 ratio (Spike: ACE2), and seeded in 96 well plates. After 48h of transfection, cells were counterstained with Hoechst. Cells were fixed with 4% paraformaldehyde for 20 minutes at room temperature and washed thrice with PBS to remove PFA and taken for imaging using Thermo Scientific CellInsight CX7 High-Content Screening (HCS) Platform. To calculate spike mutation’s effect on syncytium formation, five different fields were randomly chosen, and the area of fused cells was measured using ImageJ software.

### Western blotting

The expression of spike protein mutants was checked from the virus-producing HEK293T cells. After 48h of transfection, cells were collected and lysed in RIPA buffer supplemented with 2× PIC (protease inhibitor cocktail) and 50mM TCEP at 4°C for 30 minutes. Further, the supernatant was collected after centrifugation at 17000g for 10 minutes at 4°C. The viral particle lysate and cell lysate were run on the 8% Tris-Tricine PAGE gel following mixing lysates with 4X Laemmli. Thereafter, proteins were transferred onto the PVDF membrane (Immobilon-FL, Merck-Millipore). After the electrotransfer, the membrane was blocked with the membrane blocking reagents (Sigma) followed by primary and secondary antibody incubations for one hour at room temperature, each of which was followed by three washes with TBST. For the p24 and beta-actin detection anti p24 (NIH ARP), rabbit anti-beta actin (LI-COR Biosciences, Cat# 926-42210, RRID:AB_1850027), for SARS CoV-2 spike detection mouse anti-spike (Cat# ZMS1076, Sigma Aldrich) respectively, were used as primary antibodies. The IR dye 680 goat anti-mouse was used as a secondary antibody for the anti-spike antibody. The IR dye 800 goat anti-mouse, or IR dye 800 goat anti-rabbit (LI-COR Biosciences Cat# 925-68070, RRID:AB_2651128, and LI-COR Biosciences Cat# 925-32211, RRID:AB_2651127) were used against primary p24 and beta-actin antibody respectively.

### Software and Statistical Analysis

All graphs were generated using GraphPad Prism (version 9.0). Statistical analysis was carried out using the in-built algorithms bundled with the software. Specific portions of images were produced using Biorender. Pymol (version 1.2) and Alpha-fold were used for protein structure visualization. Western blot images were processed using Image Studio Lite Ver. 5.2 (LiCOR Biosciences). Image-J was used for image processing. The images captured using the CX7 High-content screening platform were analyzed using the Thermo Scientific HCS Studio.

## Acknowledgment

This work was supported by intramural funds from IISER Bhopal. AC is a recipient of the Wellcome Trust/DBT India Alliance Fellowship [grant number IA/I/18/2/504006]. A fellowship from the MHRD supports TM and GJ. RD is supported by a fellowship from CSIR. The authors thank Siva Umapathy for helpful discussions and Vipin Bhardwaj and Pavitra Ramdas for the technical assistance. Authors are grateful to Nevan Krogan, Massimo Pizzato, Jeremy Luban, Sonja Best, Raffaele De Francesco, Didier Trono, and the NIH AIDS Reagent Program for various reagents and cell lines.

## Declaration of interest

The authors declare no competing interests.

## Notes

### Competing Interest Statement

The authors have declared no competing interest.

## References

1. WHO. WHO Coronavirus (COVID-19) Dashboard | WHO Coronavirus (COVID-19) Dashboard With Vaccination Data. https://covid19.who.int/.

2. Rambaut, A. et al. A dynamic nomenclature proposal for SARS-CoV-2 lineages to assist genomic epidemiology. Nature Microbiology 2020 5:11 5, 1403–1407 (2020).

3. WHO. Tracking SARS-CoV-2 variants. 2021 https://www.who.int/en/activities/tracking-SARS-CoV-2-variants/.

4. Mlcochova, P. et al. SARS-CoV-2 B.1.617.2 Delta variant emergence, replication and sensitivity to neutralising antibodies. bioRxiv (2021) doi:10.1101/2021.05.08.443253.

5. Ujjainiya, R. et al. High failure rate of ChAdOx1-nCoV19 immunization against asymptomatic infection in healthcare workers during a Delta variant surge: a case for continued use of masks post-vaccination. medRxiv 2021.02.28.21252621 (2021) doi:10.1101/2021.02.28.21252621.

6. Hoffmann, M. et al. SARS-CoV-2 Cell Entry Depends on ACE2 and TMPRSS2 and Is Blocked by a Clinically Proven Protease Inhibitor. Cell 181, 271–280.e8 (2020).

7. Ferreira, I. A. T. M. et al. SARS-CoV-2 B.1.617 Mutations L452R and E484Q Are Not Synergistic for Antibody Evasion. The Journal of Infectious Diseases 224, 989 (2021).

8. Michael Rajah, M. et al. B.1.1.7 and B.1.351 SARS-CoV-2 variants display enhanced Spike-mediated fusion. bioRxiv (2021) doi:10.1101/2021.06.11.448011.

9. Liu, J. et al. BNT162b2-elicited neutralization of B.1.617 and other SARS-CoV-2 variants. Nature doi:10.1038/s41586-021.

10. Voysey, M. et al. Safety and efficacy of the ChAdOx1 nCoV-19 vaccine (AZD1222) against SARS-CoV-2: an interim analysis of four randomised controlled trials in Brazil, South Africa, and the UK. The Lancet 397, 99–111 (2021).

11. Polack, F. P. et al. Safety and Efficacy of the BNT162b2 mRNA Covid-19 Vaccine. https://doi.org/10.1056/NEJMoa2034577 383, 2603–2615 (2020).

12. Peacock, T. The SARS-CoV-2 variantsassociated with 1 infections in India, B.1.617, show enhanced spike cleavage by furin. bioRxiv (2021) doi:10.1101/2021.05.28.446163.

13. Kang, L. et al. In brief A selective sweep in the Spike gene has driven SARS-CoV-2 human adaptation. Cell 184, (2021).

14. Gupta, K. et al. Incidence of SARS-CoV-2 Infection in Health Care Workers After a Single Dose of mRNA-1273 Vaccine. JAMA Network Open 4, e2116416–e2116416 (2021).

15. Shu, Y. & McCauley, J. GISAID: Global initiative on sharing all influenza data – from vision to reality. Eurosurveillance 22, 30494 (2017).

16. Elbe, S. & Buckland-Merrett, G. Data, disease and diplomacy: GISAID's innovative contribution to global health. Global Challenges 1, 33–46 (2017).

17. Abdel Latif, A. et al. S:E156G, S:del157-158 Variant Report. 2021 https://outbreak.info/situation-reports?pango&muts=S%3AE156G&muts=S%3Adel157%2F158&loc=USA&loc=IND&loc=RUS&selected=Worldwide&overlay=false.

18. Tunyasuvunakool, K. et al. Highly accurate protein structure prediction for the human proteome. Nature 2021 596:7873 596, 590–596 (2021).

19. Mishra, T. et al. SARS CoV-2 Nucleoprotein Enhances the Infectivity of Lentiviral Spike Particles. Frontiers in Cellular and Infection Microbiology 11, 341 (2021).

20. Cele, S. et al. Escape of SARS-CoV-2 501Y.V2 from neutralization by convalescent plasma. Nature 2021 593:7857 593, 142–146 (2021).

21. Motozono, C. et al. SARS-CoV-2 spike L452R variant evades cellular immunity and increases infectivity. Cell Host & Microbe 29, 1124–1136.e11 (2021).

22. Buchrieser, J. et al. Syncytia formation by SARS-CoV-2-infected cells. The EMBO Journal 39, e106267 (2020).

23. Planas, D. et al. Reduced sensitivity of SARS-CoV-2 variant Delta to antibody neutralization. Nature (2021) doi:10.1038/s41586-021-03777-9.

24. Bussani, R. et al. Persistence of viral RNA, pneumocyte syncytia and thrombosis are hallmarks of advanced COVID-19 pathology. EBioMedicine 61, (2020).

25. Wang, P. et al. Antibody resistance of SARS-CoV-2 variants B.1.351 and B.1.1.7. Nature 2021 593:7857 593, 130–135 (2021).

26. Mlcochova, P. et al. SARS-CoV-2 B.1.617.2 Delta variant replication and immune evasion. Nature doi:10.1038/s41586-021.

27. Acevedo, M. L. et al. Infectivity and immune escape of the new SARS-CoV-2 variant of interest Lambda. medRxiv 2021.06.28.21259673 (2021) doi:10.1101/2021.06.28.21259673.

28. Kimura, I. et al. SARS-CoV-2 Lambda variant exhibits higher infectivity and immune resistance. bioRxiv 7, 2021.07.28.454085 (2021).

29. Asarnow, D. et al. Structural insight into SARS-CoV-2 neutralizing antibodies and modulation of syncytia. Cell 184, 3192–3204.e16 (2021).

30. Meng, B. et al. Recurrent emergence of SARS-CoV-2 spike deletion H69/V70 and its role in the Alpha variant B.1.1.7. Cell Reports 35, 109292 (2021).

31. Xiangyang Chi1, R. Y. J. Z.* A neutralizing human antibody binds to the N-terminal domain of the Spike protein of SARS-CoV-2. Science (New York, N.Y.) 369, 650–655 (2020).

32. Suryadevara et al. Neutralizing and protective human monoclonal antibodies recognizing the N-terminal domain of the SARS-CoV-2 spike protein. Cell 184, 2316–2331.e15 (2021).

33. McCallum Matthew et al. N-terminal domain antigenic mapping reveals a site of vulnerability for SARS-CoV-2. Cell 184, 2332–2347.e16 (2021).

34. Wibmer, C. K. et al. SARS-CoV-2 501Y.V2 escapes neutralization by South African COVID-19 donor plasma. doi:10.1101/2021.01.18.427166.

35. Gabriele, C. et al. Potent SARS-CoV-2 neutralizing antibodies directed against spike N-terminal domain target a single supersite. Cell Host & Microbe 29, 819–833.e7 (2021).

36. Harvey, W. T. et al. SARS-CoV-2 variants, spike mutations and immune escape. Nature Reviews Microbiology 2021 19:7 19, 409–424 (2021).

37. Pizzato, M. et al. A one-step {SYBR} {Green} {I}-based product-enhanced reverse transcriptase assay for the quantitation of retroviruses in cell culture supernatants. Journal of Virological Methods 156, 1–7 (2009).

38. Ramdas, P., Bhardwaj, V., Singh, A., Vijay, N. & Chande, A. Coelacanth SERINC2 Inhibits HIV-1 Infectivity and Is Counteracted by Envelope Glycoprotein from Foamy Virus. Journal of Virology 95, (2021).

39. Rosa, A. et al. HIV-1 Nef promotes infection by excluding SERINC5 from virion incorporation. Nature 526, 212–217 (2015).

40. Chande, A. et al. S2 from equine infectious anemia virus is an infectivity factor which counteracts the retroviral inhibitors {SERINC5} and {SERINC3}. Proceedings of the National Academy of Sciences 113, 13197–13202 (2016).

